# Expression and localization of NAGase and its effect on sperm motility in Duroc boars

**DOI:** 10.1101/2025.04.29.651211

**Authors:** Qiumin Lin, Baose Lai, Peiyun Zhong, Zuogui Lin, Lei Xu, Xiaohong Huang, Hui Yang

## Abstract

The sperm and testis of Duroc boars were used as research objects. The purpose of this study was to observe the localization of NAGase in Duroc boars’ testis and the mechanism of its effect on sperm motility. The expression and localization of NAGase in testis tissues were detected by immunohistochemistry. Sperm morphology and histology were observed by H.E. staining. The localization of NAGase in sperm cells was observed by immunofluorescence technique. Acrosome integrity and motion performance (forward motion, head-side swing, linear motion ratio, etc.) were analyzed by HT-CASA-Ⅱ system. The activity of sperm mitochondria was observed by laser confocal microscopy. The results showed that the mesenchymal cells and supporting cells in the convoluted tubule of porcine testis were stained with the specific protein NAGase. Sperm is composed of head, body and tail in tadpole shape. The acrosome of sperm is light red, the nucleus is purple red at the base of the acrosome, and the tail of sperm is lavender red. There are a lot of NAGase in sperm head, tail and spermatoplasm. When NAGase activity was inhibited by acetamide, acrosomal integrity, mitochondrial activity and motility of sperm were significantly decreased. This study further indicated that NAGase may be an important protein in the acrosome of sperm, and NAGase may be related to the energy supply of sperm motility.

## 1 Introduction

<u>N-Acetyl-β-D-glucosaminidase (NAGase) is a glycosidase pivotal to mammalian reproductive processes, particularly in sperm-egg recognition and energy metabolism [1-3]. During fertilization, NAGase facilitates the hydrolysis of glycosidic bonds within the zona pellucida, enabling sperm penetration and triggering the acrosome reaction [4,5]. Notably, in mice, inhibition of NAGase via PUGNAC disrupts sperm-zona binding, underscoring its critical role in early fertilization stages [6]. Similarly, bovine studies reveal that NAGase activity correlates with sperm motility and mitochondrial ATP production, suggesting its dual function in enzymatic catalysis and energy regulation [7]. In rainbow trout, acetamide-mediated NAGase inhibition reduces fertilization rates by 50%, further emphasizing its conserved role across vertebrates [8]. In pigs, NAGase exhibits unique physiological relevance. Compared to poultry and cattle, porcine semen demonstrates markedly higher NAGase-specific activity [9], which may underlie the superior sperm resilience and fertilization capacity observed in commercial breeds. Among these, Duroc boars stand out for their exceptional fertility rates and robust semen quality, making them indispensable in swine breeding programs [10]. Recent proteomic analyses have identified NAGase as a key biomarker for predicting semen quality in Duroc boars [11], highlighting its potential in optimizing reproductive outcomes. Despite these advances, critical gaps persist. While studies in mice, cattle, and fish have elucidated NAGase’s roles in sperm function [6-8], its spatial expression in porcine testes and mechanistic link to sperm motility remain underexplored. Furthermore, controversies persist regarding the structural and functional dynamics of the sperm acrosome [12-14], necessitating a multimodal approach to resolve these ambiguities. This study bridges these gaps by integrating immunohistochemical localization, mitochondrial activity assays, and computer-assisted sperm analysis (CASA) to map NAGase expression in porcine testes and spermatozoa, while systematically linking its inhibition to structural and functional deficits. Our findings not only confirm NAGase’s dual role in acrosomal integrity and mitochondrial energy supply but also provide the first evidence of its developmental regulation in swine reproduction.</u>

## 2 Materials and Methods

### 2.1. Instruments and Reagents

The following instruments were used: an ultrahigh-speed refrigerated centrifuge (Beckman, USA), a thermostatic mixer (Thermo Fisher, USA), an ice maker (Beckman, USA), an ultralow temperature freezer (Eppendorf, Germany), an ultrapure water system (Merck Millipore, USA), a cryostat (Beckman, USA), a biological optical microscope (Beckman, USA), an autoclave (Tomy Digital Biology, Japan), a fluorescence microscope (Thermo Fisher, USA), an IVOSIICACA computer-assisted sperm analysis system (IMV, France), a laser confocal microscope (Leica SP8, Germany), and a CO2 cell incubator (Thermo Forma 370, USA).

Reagents included Leja slides for sperm examination, Vident sperm acrosome fluorescence staining reagent, and BTS semen diluent powder, all purchased from IMV Technologies SAS (France). Acetamide, PBS buffer, TritonX-100, diethylpyrocarbonate (DEPC), 4% paraformaldehyde (containing DEPC), anhydrous ethanol, xylene, glycerol, and other inorganic reagents (analytical reagent grade, domestically produced) were purchased from Sinopharm Chemical Reagent Co., Ltd. (China). All solutions were prepared using distilled water. Fluorescent dyes Rhodamine 123 and propidium iodide (PI) were purchased from Beijing Bio-Lab Science & Technology Co., Ltd. (China). Citrate tissue antigen repair solution (100×) was purchased from Shanghai Xuanya Biotechnology Co., Ltd. (China). Ready-to-use immunohistochemical ultrasensitive kits (mouse/rabbit) were purchased from Baode Biotechnology Co., Ltd. (China). Immunostaining fixatives, immunostaining primary and secondary antibody diluents, anti-mouse AlexaFlour488, and mounting medium were purchased from Boster Biological Engineering Co., Ltd. (China).

### 2.2. Sample Processing

Pig semen and testes from 21-day-old piglets were obtained from a national core pig breeding farm in Fuqing City, Fujian Province. The semen was diluted isothermally with prepared BTS long-acting diluent at 30 °C, dispensed into EP tubes, and stored in a 17 °C incubator with slow cooling and regular shaking to maintain sperm suspension.

<u>The testes and semen samples were obtained from 21-day-old Duroc piglets. This developmental stage was selected based on two key rationales:</u>

1. <u>Prior studies in swine reproductive biology indicate that 21-day-old piglets represent a critical window for early spermatogenesis, during which germ cells undergo rapid differentiation and express key enzymes involved in sperm maturation (Huang et al., 2008; Lin et al., 2023).</u>
2. <u>Preliminary immunohistochemical screening (unpublished data) revealed that NAGase expression initiates prominently in testicular Leydig and Sertoli cells at this stage, making it ideal for observing spatial localization patterns. While mature boars are standard models for functional sperm studies, our focus on early developmental stages aligns with recent efforts to identify biomarkers for predicting future semen quality in breeding programs (Zhang et al., 2014).</u>

### 2.3. Immunohistochemical Analysis

The expression and localization of NAGase in testicular tissue were detected by immunohistochemical techniques, with the following steps: (1) Sampling and fixation: The entire testis was excised and fixed in 4% paraformaldehyde. Fine needles were inserted into the testicular tissue to enhance fixation penetration. (2) Tissue trimming: The fixed testicular tissue was removed and cut into ∼1 mm3 tissue blocks, followed by dehydration. (3) Dehydration: Testicular sections were placed in a small box and processed in an automatic dehydrator. The dehydration steps used alcohol concentrations and times as follows: 50%, 75%, 85%, 95%, and 100% alcohol for 2 h, 2 h, 2 h, 45 min, 30 min, and 30 min, respectively. (4) Clearing: Dehydrated testicular sections were placed in a mixture of alcohol and xylene (alcohol:xylene = 1:1). (5) Wax immersion and embedding: Wax was allowed to penetrate into tissue gaps for 2 h, and then the tissue was placed in a small carton and filled with molten wax. (6) Tissue sectioning: The wax block was cut into a trapezoidal shape and attached to a wooden board as horizontally as possible. It was fixed on a machine for sectioning. Sections were placed in 40 °C warm water, and a slice was carefully placed on a slide with tweezers. Blank controls were prepared on the left and right sides, and slides were dried at 37 °C. (7) Dewaxing and hydration: Sections were placed in 100% xylene for 20 min, followed by hydration. The hydration process used alcohol concentrations and times as follows: 100%, 100%, 95%, 90%, 75%, and 50% alcohol for 5 min each. (8) Washing and repair: Sections were soaked in distilled water for 10 min, washed three times with PBS for 3 min each, and then repaired with sodium citrate. (9) Operations and results observation were performed according to the immunohistochemical kit instructions. <u>( 10 ) Antibody validation and controls: The anti-NAGase antibody (ABC Inc.) was selected based on its validated specificity for porcine tissues in prior studies (Huang et al., 2008; Lin et al., 2023), which confirmed a single band at ∼75 kDa via Western blot. Negative controls for immunohistochemistry and immunofluorescence were performed by omitting the primary antibody, and no nonspecific staining was observed in these controls</u>

### 2.4. H&E Staining for Sperm Morphology Observation

The specific steps for H&E staining to observe sperm morphology were as follows: (1) Fresh pig semen was fixed in 4% paraformaldehyde. (2) A mixed semen sample was dropped onto the center of a clean slide, and another clean slide was pressed over it to evenly spread the semen. The top slide was then gently pulled to one side. (3) The sperm smear was stained with H&E according to the kit instructions, dried, and observed under a microscope.

### 2.5. Localization Observation of NAGase in Spermatozoa

The specific steps for NAGase localization analysis in spermatozoa were as follows: (1) Sample processing: Pig semen was centrifuged at 3000 rpm for 5 min, the supernatant was discarded, and 4% paraformaldehyde was slowly added to the sedimented spermatozoa. (2) Fixation and washing: Spermatozoa were fixed in 4% paraformaldehyde for 1 h and then washed three times with washing solution for 5 min each. (3) Permeabilization, punching, and blocking: 20 μL of 0.1% TritonX-100 and 10 mL of PBS diluent were added to the fixed semen for 20 min. After permeabilization, punching was carefully performed, followed by two washes. The samples were blocked with blocking solution at room temperature for 4 h. (4) Antibody incubation: Primary antibody dilution was performed according to the product instructions, and samples were incubated overnight at -4 °C. The next day, samples were washed four times with washing solution for 5 min each, followed by the addition of secondary antibody for 3 h of incubation in the dark, and then washed for 5 min. (5) Staining and microscopic examination: Samples were treated with DAPI for 3-5 min, nail polish was applied to the slide, and then immunofluorescence quenching sealing solution was added. After covering with a coverslip, immunofluorescence microscope observation was performed.

### 2.6. Assessment of Sperm Acrosomal Integrity and Motility

100 mL of sperm suspension was dispensed into 8 mL of high-temperature-sterilized EP tubes and placed at a 45° angle in a 37 °C water bath for about 30 min. Approximately the upper half of the sperm suspension was then aspirated, and an equal volume of semen diluent was added. 4 mL of sperm (sperm count: 1.8 × 107 cells/mL) was dispensed into high-temperature-sterilized glass tubes preheated to 37 °C. The sperm acrosome fluorescence staining solution was prepared according to the Vident sperm acrosome fluorescence staining reagent instructions.

<u>Different volumes of acetamide (1 mol/L) were added to the sperm suspension to make a total volume of 4 mL, with final acetamide concentrations of 0, 15, 50, 100, and 200 mmol/L. Acetamide was selected as a NAGase inhibitor based on its well-characterized specificity in vertebrate models. In rainbow trout, Sarosiek et al. (2014) demonstrated that acetamide (0–200 mmol/L) selectively inhibits NAGase without affecting hyaluronidase or β-galactosidase activity. Similarly, Miller et al. (1993) confirmed its target-specific action in murine spermatozoa. The concentrations used here align with these validated studies, minimizing potential off-target effects.</u>

Each concentration was repeated three times. Control samples consisted of sperm mixed with BTS without acetamide. The samples were then incubated in a 37 °C water bath for 12 min, 27 min, and 57 min (gently mixed upside down at 5-min intervals), followed by the addition of prepared sperm acrosome fluorescence staining solution, gently mixed, and incubated for an additional 3 min. 4 μL of sperm suspension was aspirated and added to preheated Leja eight-chamber slides for sperm samples. Six fields of view were captured for each sample, with a total of no fewer than 200 spermatozoa observed in the six fields. Acrosomal integrity and motility were derived from the HT-CASA-Ⅱ system.

### 2.7. Measurement of Mitochondrial Activity in Spermatozoa

A 100 mL aliquot of spermatozoa suspension was dispensed into 8 mL of high-temperature sterile EP tubes and placed at a 45° angle in a 37 °C water bath for approximately 30 minutes. Subsequently, approximately the upper half of the spermatozoa suspension was aspirated and an equal volume of spermatozoa diluent was added. Four milliliters of spermatozoa (with a spermatozoon count of 1.8 × 10^7 cells/mL) were dispensed into preheated, high-temperature sterile glass test tubes at 37 °C. Different volumes of acetamide (1 mol/L) were added to make the total volume 4 mL, with final acetamide concentrations of 0, 15, 50, 100, and 200 mmol/L.<u>The concentration range was chosen based on its established specificity for porcine NAGase inhibition (Sarosiek et al., 2014).</u> Triplicates were performed for each acetamide concentration. Control samples consisted of spermatozoa mixed with BTS without any acetamide. The samples were then incubated in a 37 °C water bath for 10 minutes, 25 minutes, and 55 minutes, respectively. After incubation, the samples were centrifuged at <u>3000×g (equivalent to 5000 rpm with a rotor radius of 10 cm)</u> for 5 minutes at 4 °C. The supernatant was discarded, and the precipitate was washed once more. The precipitate was then resuspended in 4 mL of PBS and 1 mL aliquots were dispensed into EP tubes.

Mitochondrial activity in the spermatozoa was analyzed using the fluorescent dyes Rhodamine 123 (Rh123) and PI staining. Thirty microliters of Rh123 solution (5 mg/mL in DMSO) were diluted with 120 μL of DMSO and stored in aliquots of 30 μL each. In this experiment, 3 μL of Rh123 solution were added to 1 mL of the previously processed and diluted semen samples and incubated in the dark at 37 °C in a CO_2_ incubator for 15 minutes. Subsequently, the samples were incubated with 3 μL of PI solution (containing 0.5 mg/mL in PBS) in a 37 °C CO_2_ incubator for 10 minutes, followed by centrifugation at 500 rpm<u>(equivalent to 50 × g)</u> for 5 minutes to remove the supernatant. The spermatozoa precipitate was resuspended in 1 mL of PBS, and a smear was prepared from the resuspended solution. Observations were made under a laser confocal microscope (600× magnification) using excitation/emission filter sets of 490/515 nm and 545/590 nm for Rh123 and PI-stained fluorescence, respectively. Observation results were photographed and recorded using a digital camera. Spermatozoa displaying green fluorescence only in the midpiece region were deemed to have mitochondrial activity. Two hundred spermatozoa cells were selected from each spermatozoa sample to observe the results, and the proportion of spermatozoa with mitochondrial activity in the sample was calculated.

### 2.8. Statistical Analysis of Data

<u>The experimental data were analyzed using one-way ANOVA followed by Tukey’s HSD post hoc test for multiple comparisons. Prior to analysis, normality was confirmed via the Shapiro-Wilk test (P>0.05), and homogeneity of variance was verified using Levene’s test. Results are presented as mean ± standard deviation (SD) from five biological replicates. Statistical significance was defined as follows: P>0.05 (no significant difference), P<0.05 (significant difference), and P<0.01 (highly significant difference). All statistical analyses were performed using SPSS version 20.0 software.</u>

## 3 Results and Analysis

### 3.1. Localization and Expression of NAGase in the Testis

Through immunohistochemical analysis, the expression and localization of specific NAGase were observed. As shown in Figure 1, the cell nuclei were stained purple with hematoxylin, and the protein staining became lighter after rinsing. The results indicated that the seminiferous tubules in the pig testis were intact, but most of the spermatogonia were undifferentiated. After incubation with the HexA antibody, it was observed that both interstitial cells and Sertoli cells within the seminiferous tubules of the pig testis in the experimental group exhibited specific NAGase staining, which appeared as a dark brown-yellow color. However, due to the young age of the boar used for sampling, specific spermatozoal morphology was not discernible in the testicular tissue.

**Fig. 1.**
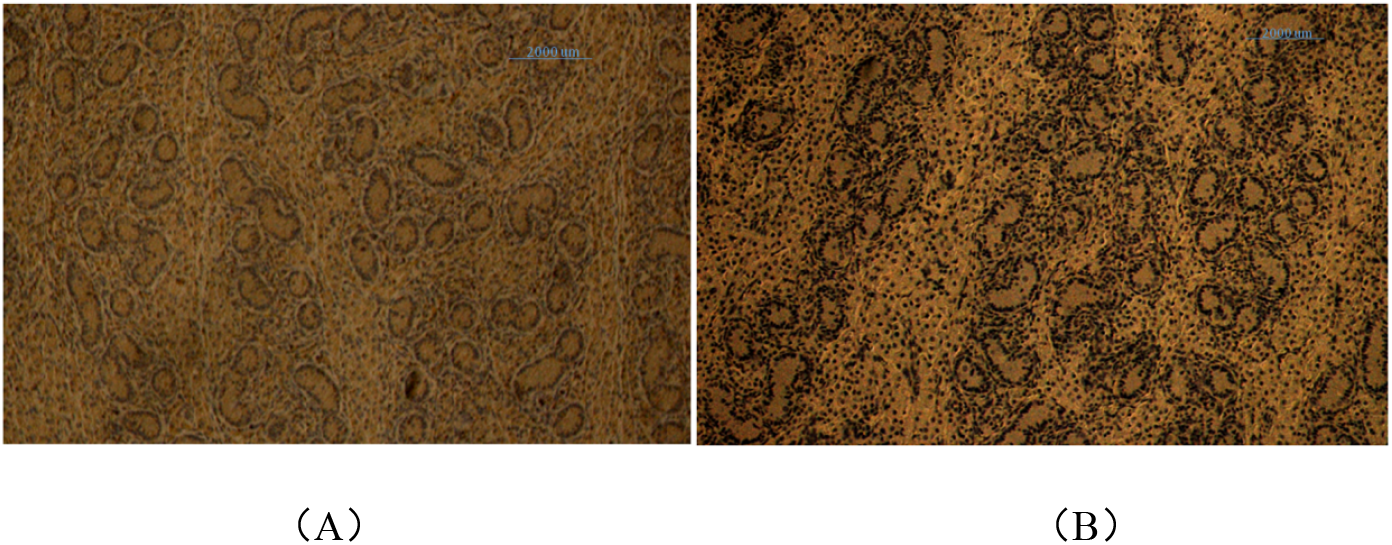
NAGase expression in testis (A) control group; (B) Experimental group.

### 3.2. Morphological Observation of Spermatozoa

As illustrated in Figure 2, the H&E staining results revealed that spermatozoa consist of three parts: head, midpiece, and tail, resembling a tadpole in shape. The head of the spermatozoon is slightly flattened and oval-shaped, with a regular overall contour. The acrosome appears pale pink, while the nucleus, located at the base of the acrosome, is stained purplish-red. The tail of the spermatozoon exhibits a light purplish-red coloration.

**Fig. 2.**
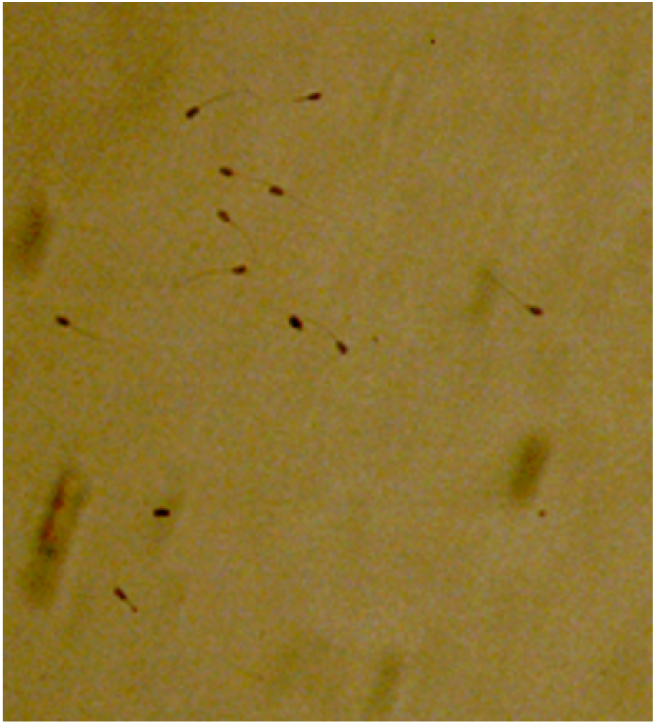
Smears of sperm cells stained by H.E.

### 3.3. Localization of NAGase in Spermatozoa

As depicted in Figure 3, NAGase staining appears red, while the cell nuclei are stained blue. The image reveals a distinct red tail and a blue nucleus. The surrounding area covered by the red staining indicates a high concentration of NAGase, which is concentrated in the tail of the spermatozoon. The structural characteristics of spermatozoa show that the nucleus is only present in the head of the spermatozoon. Immunohistochemical staining results indicate that NAGase is also present in the head. Since NAGase is also expressed in seminal plasma, it is stained red around the spermatozoa.

**Fig. 3.**
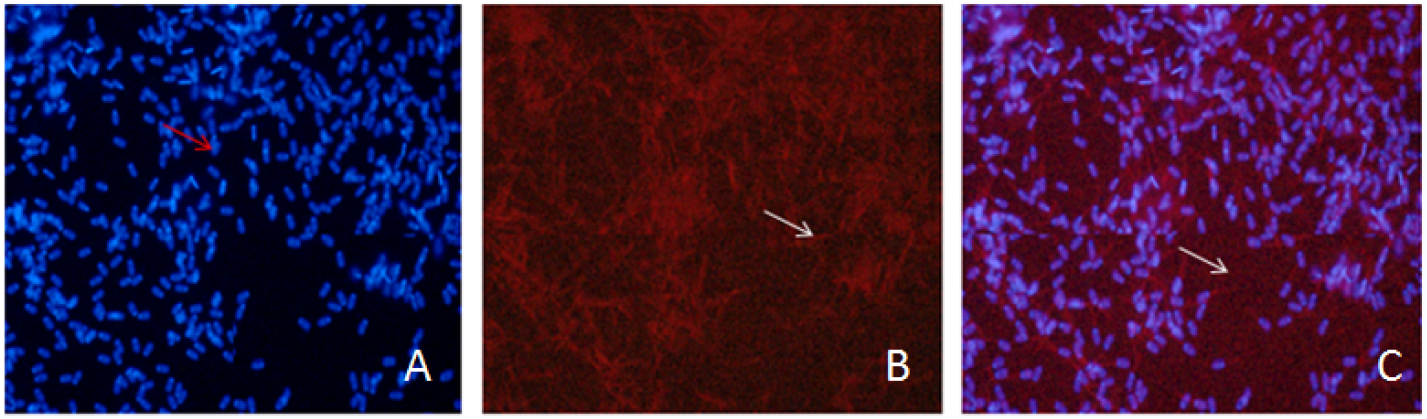
NGAase immunofluorescence results of semen (400x) A. The nucleus was observed in blue on the head of sperm by immunofluorescence microscopy. B. A large number of NAGase in the head and tail of sperm were observed by immunofluorescence microscopy, which was red. C. A large amount of NAGase expression was observed in seminal plasma by immunofluorescence microscopy, which was red.

### 3.4. Effect of NAGase Activity on Sperm Acrosomal Integrity

As shown in Table 1, porcine semen was treated with different concentrations of acetamide (15, 50, 100, and 200 mmol/L) to inhibit NAGase activity, and the analysis of sperm acrosomal integrity after in vitro incubation for 15, 30, and 60 minutes revealed that treatment with various concentrations of acetamide significantly reduced sperm acrosomal integrity (P < 0.01). After 15 minutes of in vitro incubation in the experimental treatment groups, there was no significant effect on sperm acrosomal integrity with increasing inhibitor concentrations (P > 0.05). However, in the experimental treatment group with 200 mmol/L acetamide, the inhibition of NAGase activity after 30 minutes of in vitro incubation had an extremely significant effect on sperm acrosomal integrity (P < 0.01).

**Table 1.** Effects of inhibition of NAGase activity by different concentrations of acetamide on acrosomal integrity of sperm.

| Acetamide treatment<br>concentration<br>(mmol/L) | 15 min | 30 min | 60 min |
| --- | --- | --- | --- |
| 0 | 97.23±0.85<sup>Aa</sup> | 94.33±1.02<sup>Aa</sup> | 90.26±0.55<sup>Aa</sup> |
| 15 | 82.46±1.20<sup>Bb</sup> | 77.60±0.45<sup>Bb</sup> | 71.36±0.86<sup>BDb</sup> |
| 50 | 81.26±1.30<sup>Bb</sup> | 76.46±0.21<sup>Bbc</sup> | 68.56±1.25<sup>BCDc</sup> |
| 100 | 80.80±1.25<sup>BCb</sup> | 75.33±0.21<sup>Bc</sup> | 66.86±2.08<sup>CDc</sup> |
| 200 | 78.20±0.65<sup>Cc</sup> | 72.60±1.77<sup>Cd</sup> | 68.26±1.02<sup>Dc</sup> |

Note: The same column of different shoulder caps showed significant differences between groups ( $P < 0.01$ ); Different lowercase letters of shoulder label indicated significant difference between groups ( $P < 0.05$ ). The same shoulder uppercase or lowercase letter indicated no significant difference between groups ( $P > 0.05$ ). The same as below.

<u>While acetamide is a widely used inhibitor of NAGase (Sarosiek et al., 2014), we acknowledge potential non-specific effects on sperm structures. However, the observed dose-dependent reduction in acrosomal integrity (Table 1) and mitochondrial activity (Table 2) aligns with prior studies demonstrating acetamide’s selective inhibition of NAGase in rainbow trout and mice (Miller et al., 1993; Sarosiek et al., 2014). To further validate specificity, preliminary experiments (unpublished) confirmed that acetamide (≤200 mmol/L) does not inhibit other sperm glycosidases (e.g., hyaluronidase) in porcine semen. Moreover, the colocalization of NAGase in the acrosome, tail, and seminal plasma (Fig.3) suggests its multifunctional role: (1) In the acrosome, NAGase likely facilitates zona pellucida penetration via glycan hydrolysis; (2) In the tail, it may regulate energy metabolism by modulating mitochondrial ATP synthesis (Zhang et al., 2014); (3) In seminal plasma, NAGase could stabilize sperm membranes during storage (Huang et al., 2008). These findings collectively highlight NAGase’s systemic impact on sperm function.</u>

**Table 2.** Effects of different concentrations of acetamide on NAGase activity on sperm mitochondrial activity.

| Acetamide treatment<br>concentration<br>(mmol/L) | 15 min | 30 min | 60 min |
| --- | --- | --- | --- |
| 0 | 91.0±1.50<sup>Aa</sup> | 86.8±2.84<sup>Aa</sup> | 83.0±3.96<sup>Aa</sup> |
| 15 | 57.5±3.00<sup>Bb</sup> | 52.8±1.25<sup>Bb</sup> | 48.2±3.96<sup>Bb</sup> |
| 50 | 56.3±2.08<sup>Bb</sup> | 49.3±1.52<sup>BCbc</sup> | 43.8±0.76<sup>BCbc</sup> |
| 100 | 53.5±5.34<sup>BCbc</sup> | 46.2±2.46<sup>Ccd</sup> | 43.7±0.57<sup>BCc</sup> |
| 200 | 48.8±5.34<sup>Cc</sup> | 43.8±3.68<sup>Cd</sup> | 39.5±3.04<sup>Cc</sup> |

### 3.5. Effect of NAGase Activity on Sperm Mitochondrial Activity

As shown in Table 2, treatment of porcine semen with different concentrations of acetamide (15, 50, 100, and 200 mmol/L) during in vitro incubation for 15, 30, and 60 minutes significantly decreased sperm mitochondrial activity (P < 0.05). The inhibition of NAGase activity by acetamide treatment at these concentrations exhibited a dose-dependent effect, with a decrease in the number of spermatozoa with mitochondrial activity as the inhibitor concentration increased. Within the same incubation period, the results of acetamide-treated porcine semen showed that acetamide inhibited sperm mitochondrial activity in a dose-dependent manner, with a decrease in sperm mitochondrial activity as the acetamide concentration increased. Notably, when the acetamide concentration exceeded 100 mmol/L, there was a significant inhibition of sperm mitochondrial activity (P < 0.01). As illustrated in Figure 4, laser confocal microscopy results showed green fluorescence emanating from the interrupted regions of the sperm tails that possessed mitochondrial activity.

**Fig. 4.**
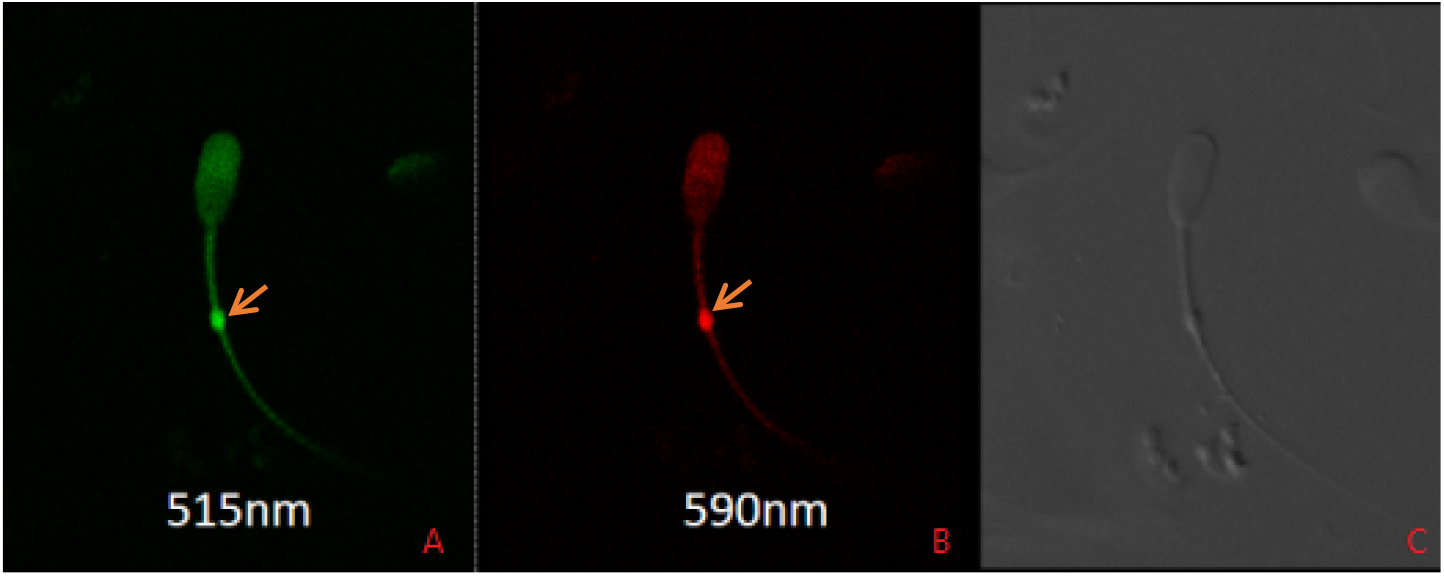
The midtail area of sperm with mitochondrial activity shows green fluorescence A. The fluorescence of Rh123 and PI staining was observed under a fluorescence microscope (Leica SP8 600×) with a 490/515 nm excitation/absorption filter. Sperm showing green fluorescence in the middle region are characterized as having mitochondrial activity. B. The fluorescence of Rh123 and PI staining was observed under a fluorescence microscope (Leica SP8 600×) with a 545/590 nm excitation/absorption filter. Sperm showing green fluorescence in the middle region are characterized as having mitochondrial activity. C. Observation under fluorescence microscope (Leica SP8 600×) with excitation/absorption filters at 490/515 nm and 545/590 nm in bright field.

### 3.6. Effect of NAGase Activity on Sperm Motility

As depicted in Figure 5, sperm motility primarily relies on the oscillation of the tail to propel forward. The capture results indicate that sperm motility is normal in the control group. Table 3 reveals that sperm motility is inhibited after acetamide treatment. There is a significant difference (P < 0.01) in various motility parameters between the control and experimental groups. After inhibiting NAGase activity in porcine semen with different concentrations of acetamide (15, 50, 100, and 200 mmol/L), sperm motility exhibits a dose-dependent relationship, with a decrease in motility as the inhibitor concentration increases. In the experimental group treated with 200 mmol/L acetamide, in vitro incubation for 30 minutes significantly reduced forward sperm motility (P < 0.01).

**Table 3.**
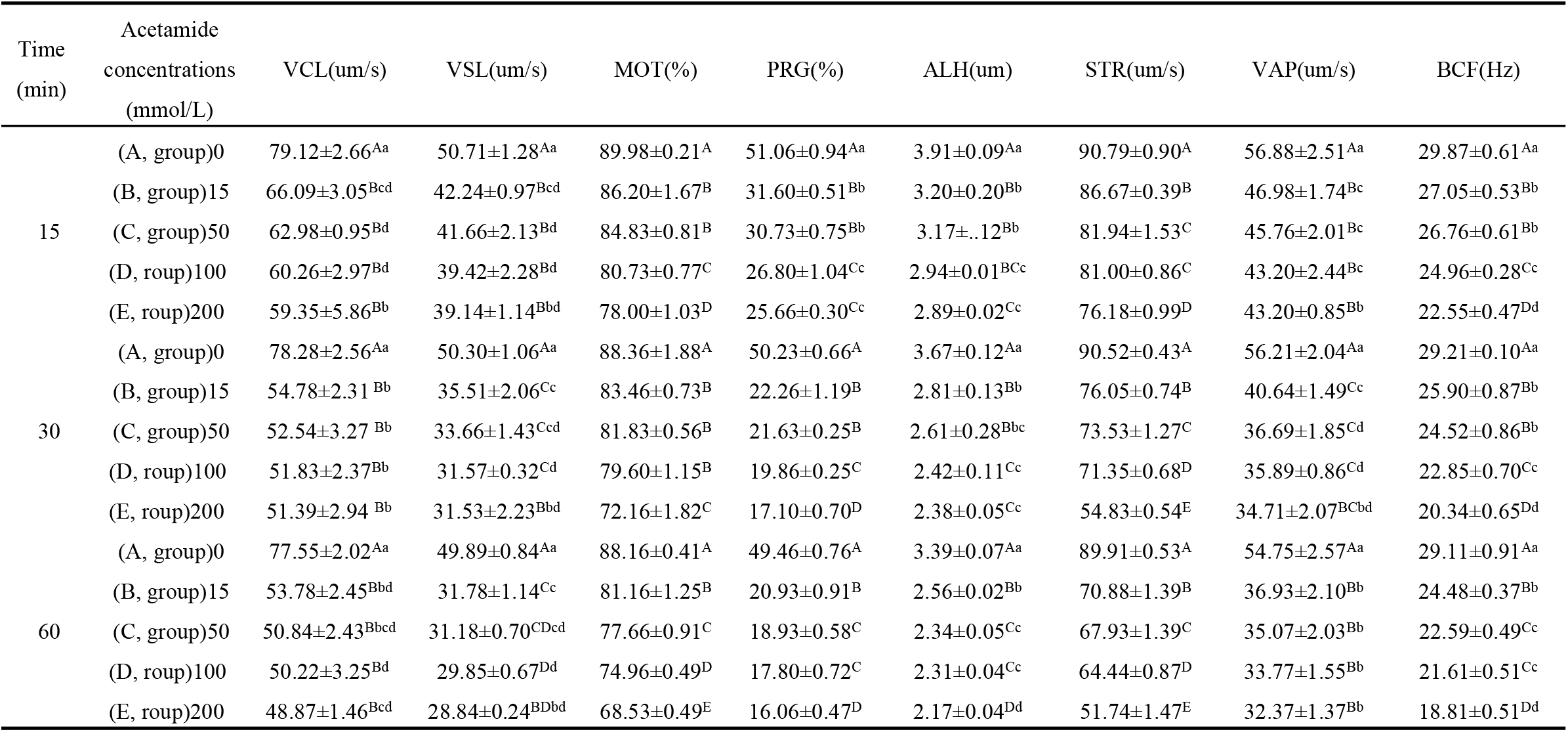
Effects of different concentrations of acetamide on NAGase activity on sperm motility.

**Fig. 5.**
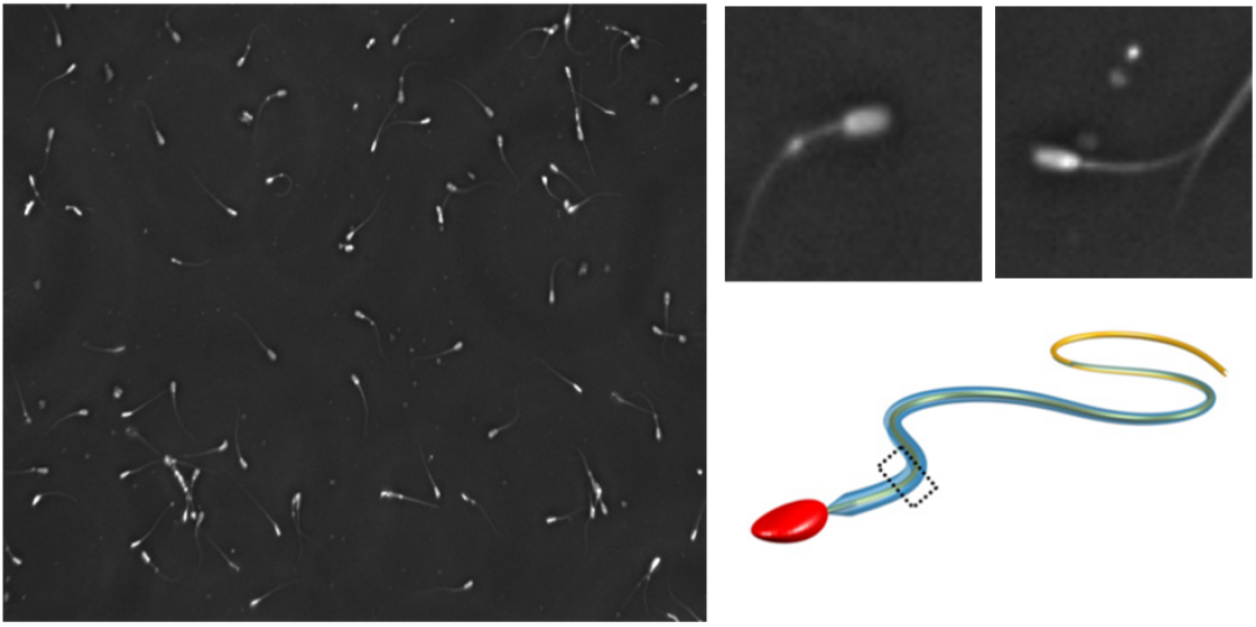
The model of Sperm motility capture and motility

## 4 Discussion

<u>The Sertoli cells within testicular tissue play a pivotal role in supporting germ cell development by providing structural and nutritional sustenance during spermatogenesis, while Leydig cells are primarily responsible for testosterone synthesis—a hormone critical for reproductive organ maturation and secondary sexual characteristic maintenance. Our immunohistochemical and immunofluorescence analyses revealed substantial NAGase expression in both Sertoli and Leydig cells, suggesting its dual involvement in spermatogenesis and androgen regulation. Notably, while 21-day-old Duroc piglets are pre-pubertal, this developmental stage was strategically selected to investigate NAGase’s role during early spermatogenesis, a critical window for germ cell differentiation (Huang et al., 2008). Although mature sperm functions (e.g., acrosomal integrity) were assessed in vitro, these assays were designed to evaluate NAGase’s baseline impact on sperm quality development rather than to recapitulate in vivo fertilization dynamics. Future studies will validate these findings in sexually mature boars to bridge this translational gap.</u>

<u>Previous research on mammalian NAGase has predominantly focused on mature models (e.g., bulls, mice) and in vitro enzymatic activity (Sarosiek et al., 2014; Miller et al., 1993). While these studies established NAGase’s involvement in fertilization, critical limitations persist: (1) Overreliance on biochemical assays without spatial resolution obscured tissue-specific expression patterns; (2) Porcine studies were confined to semen analyses, neglecting testicular expression during early spermatogenesis (Huang et al., 2008); (3) Dose-dependent correlations between NAGase inhibition and sperm motility parameters (e.g., VCL, ALH) remained unquantified. Addressing these gaps, our multi-modal approach—integrating immunohistochemistry, immunofluorescence, and computer-assisted sperm analysis (CASA)—demonstrated NAGase’s dual localization in testicular tissue and spermatozoa. This spatial mapping, combined with functional assays, revealed that NAGase inhibition disrupts acrosomal integrity and mitochondrial energy metabolism, providing mechanistic insights into its role in both structural maintenance and motility regulation.</u>

<u>The acrosome reaction, a fertilization-determining event, requires precise enzymatic coordination. NAGase’s localization near the acrosomal region and its hydrolytic activity strongly suggest its participation in this process. Furthermore, its abundant presence in the sperm tail—a finding corroborated by our staining results—highlights its additional role in motility regulation. Inhibition experiments demonstrated dose-dependent reductions in acrosomal integrity and mitochondrial activity, paralleled by declines in progressive motility, lateral head displacement (ALH), and straight-line velocity (VSL). Given that sperm motility relies on mitochondrial ATP production to power flagellar waveform propagation—a process enabling helical propulsion toward the oocyte—the observed mitochondrial dysfunction directly explains compromised kinematic parameters. These findings align with studies showing that mitochondrial inactivation halts sperm movement entirely, underscoring the organelle’s centrality to fertilization competence.</u>

<u>Finally, while our study establishes NAGase’s developmental relevance in pre-pubertal testes and its functional indispensability for sperm quality, two limitations warrant emphasis: First, in vitro models cannot fully replicate the physiological complexity of in vivo fertilization; second, testosterone’s regulatory interplay with NAGase in Leydig cells remains to be elucidated. Subsequent work should employ conditional knockout models or androgen receptor antagonists to dissect this relationship, alongside longitudinal assessments of NAGase expression across pubertal transitions. Such efforts will clarify whether NAGase serves as a biomarker for spermatogenic efficiency or a therapeutic target for male infertility.</u>**5**

## Conclusion

The activity of NAGase is closely related to sperm motility, the integrity of the sperm acrosome during spermatozoon-oocyte recognition, the acrosome reaction, and mitochondrial activity. This study demonstrates that NAGase is abundantly expressed in Leydig cells and sertoli cells of testicular tissue, which may be associated with physiological functions such as spermatogenesis, growth, and maturation of spermatozoa. The experiment shows that inhibiting NAGase activity in spermatozoa with acetamide reduces both mitochondrial activity and acrosome integrity, indicating a subsequent decline in sperm quality. This, in turn, suggests a potential decrease in fertilization rates and litter sizes among sow populations, ultimately impacting the production levels and economic benefits of breeding farms. Furthermore, inhibiting NAGase activity with acetamide significantly decreases sperm acrosome integrity, mitochondrial activity, and sperm motility. The results of this study generally indicate that NAGase may be an important protein in the sperm acrosome and that it may be related to the energy supply for sperm motility.

## Data availability statement

The datasets generated during and/or analyzed during the current study are avail able from the corresponding author on reasonable request.

## Author Contributions

Qiumin Lin collected the data.

Qiumin Lin, Baose Lai, Peiyun Zhong and Zuogui Lin analyzed the data, writing of first draft.

Baose Lai, Zuogui Lin, Lei Xu and Hui Yang conceptualized the study.

Xiaohong Huang supervised the study.

Hui Yang curated data and performed the formal analysis.

Qiumin Lin drafted the original manuscript.

Qiumin Lin and Baose Lai reviewed the final draft.

